# Scaffolding and Completing Genome Assemblies in Real-time with Nanopore Sequencing

**DOI:** 10.1101/054783

**Authors:** Minh Duc Cao, Son Hoang Nguyen, Devika Ganesamoorthy, Alysha G. Elliott, Matthew Cooper, Lachlan J.M. Coin

**Affiliations:** Institute for Molecular Bioscience, the University of Queensland, St Lucia, Brisbane, QLD 4072 Australia; Department of Genomics of Common Disease, Imperial College London, London W12 0NN, UK

## Abstract

Genome assemblies obtained from short read sequencing technologies are often fragmented into many contigs because of the abundance of repetitive sequences. Long read sequencing technologies allow the generation of reads spanning most repeat sequences, providing the opportunity to complete these genome assemblies. However, substantial amounts of sequence data and computational resources are required to overcome the high per-base error rate inherent to these technologies. Furthermore, most existing methods only assemble the genomes after sequencing has completed which could result in either generation of more sequence data at greater cost than required or a low-quality assembly if insufficient data are generated. Here we present the first computational method which utilises real-time nanopore sequencing to scaffold and complete short-read assemblies while the long read sequence data is being generated. The method reports the progress of completing the assembly in real-time so users can terminate the sequencing once an assembly of sufficient quality and completeness is obtained. We use our method to complete four bacterial genomes and one eukaryotic genome, and show that it is able to construct more complete and more accurate assemblies, and at the same time, requires less sequencing data and computational resources than existing pipelines. We also demonstrate that the method can facilitate real-time analyses of positional information such as identification of bacterial genes encoded in plasmids and pathogenicity islands.

## Introduction

High-throughput sequencing technology has profoundly transformed genomics research over the last decade with the ability to sequence the whole genome of virtually every organism on the planet. Most sequencing projects to date employ short read technology and hence cannot unambiguously resolve repetitive sequences which are present abundantly in most genomes. As a result, the assemblies are fragmented into large numbers of contigs and the positions of repeat sequences in the genome cannot be determined. These repeat sequences often play important biological roles. For example, they mediate lateral transfer of pathogenicity islands between bacterial species. Analysing these regions is thus essential for determining key characteristics such as antibiotic resistance profiles and for identifying highly pathogenic variants of many bacterial species (Ashton et al., 2015).

Long read sequencing technologies introduced recently by Pacific Biosciences (SMRT sequencing) and Oxford Nanopore (nanopore sequencing) permit the generation of reads spanning most repetitive sequences which can be used to close gaps in the fragmented assemblies. The key innovation of the MinION nanopore sequencing device is that it measures the changes in electrical current as a single-stranded DNA passes through the nanopore and uses the signal to determine the nucleotide sequence of the DNA strand (Branton et al., 2008;Kasianowicz et al., 1996 Kasianowicz et al., 1996; Stoddart et al., 2009). As such the raw data of a read can be retrieved and analysed while sequencing is still in progress. This offers the opportunity to obtain analysis results as soon as sufficient data are generated, upon which the sequencing can be terminated or used for other experiments.

A number of algorithms have been developed to make use of the long reads for genome assembly. *De novo* assemblers such as HGAP (Chin et al., 2013), Canu (Berlin et al., 2015) and nanocorrect/nanopolish (Loman et al., 2015) are able to completely assemble a bacterial genome using only long read sequencing data. However, because of the high error rates in these sequencing technologies, this *de novo* approach requires substantial amounts of sequencing data and extensive computational resources, mainly for polishing the genome assembly. The hybrid assembly approach, which combines error-prone long reads with highly accurate and cheaper short read sequence data, provides a more economical and efficient alternative for building complete genomes. Tools in this category generally a) error-correct long reads with the high quality short reads, and assemble the genome with the corrected long reads (PBcR (Koren et al., 2012), Nanocorr (Goodwin et al., 2015) and NaS (Madoui et al., 2015), or b) use long reads to scaffold and to fill in gaps of the assemblies from short read sequencing (SPAdes-hybrid (Ashton et al., 2015;Bankevich et al., 2012), SSPACE-LongRead (Boetzer and Pirovano, 2014;Karlsson et al., 2015) and LINKS (Warren et al., 2015).

While these tools are reported to assemble high quality bacterial genomes, they have not made use of real-time sequencing potential of the MinION; assembly of a genome can only be performed in ‘batch mode’ after the sequencing is complete. This can lead to over-sequencing, in which extra cost and time is incurred to generate an assembly which could have been generated with fewer data; or under-sequencing resulting in a low-quality assembly. Here, we present npScarf, the first hybrid assembler that can scaffold and complete fragmented short read assemblies with sequence data streaming from the MinION while sequencing is still in progress. npScarf can constantly report the quality of the assembly during the experiment so that users can terminate the sequencing when an assembly of sufficient quality and completeness is obtained. We show that npScarf can generate more accurate and more complete genomes than existing tools while requiring less nanopore sequencing data and computation resources. We also demonstrate that, npScarf can facilitate the real-time analysis of positioning genomic sequences such as identifying genes encoded in plasmids and pathogenicity islands underlying the acquisition of antibiotic resistance.

**Figure 1:**
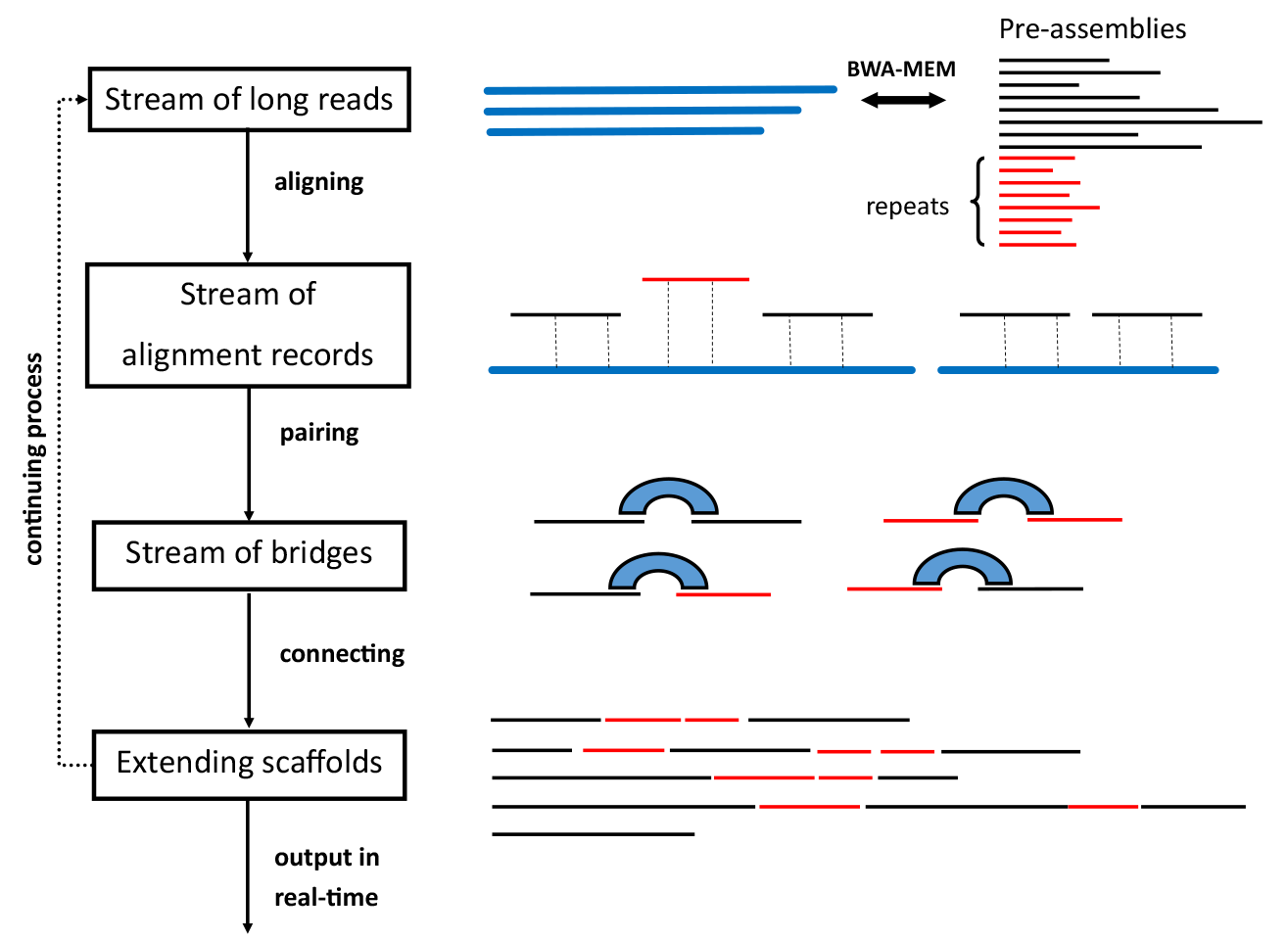
General workflow of the real-time algorithm

## Results

### Algorithm overview

The genomes of most organisms contain an abundance of repeat sequences that are longer than the read length limit (300bps) of Illumina sequencing platforms (Treangen and Salzberg, 2012). In assembling a genome using this technology, these repeat sequences cannot be distinguished and hence are often collapsed into contigs, leaving gaps in the genome assembly. To scaffold and fill in gaps in the assembly, npScarf first determines the multiplicity of each contig, thereby identifying contigs representing non-repetitive sequences (called unique contigs). These unique contigs are then bridged with long reads to form a backbone of the genome, while repetitive contigs are used to fill in gaps in the backbone.

### Determining unique contigs

Prior to scaffolding a fragmented short read genome assembly, npScarf determines the multiplicity of each contig in the assembly by comparing short read sequencing coverage of the contig to that of the whole genome. The coverage information is often included in the sequences assembled by most tools such as SPAdes (Bankevich et al., 2012) and Velvet (Zerbino and Birney, 2008), or otherwise can be obtained from mapping of short reads to the assembly. npScarf estimates the depth coverage of the genome as the normalised average coverage of up to 20 largest contigs longer than 20Kb, which are most likely unique contigs in bacterial genomes (Koren and Phillippy, 2015).

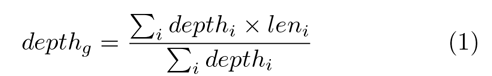

In Equation 1, *depth_i_* and *len_i_* represent the sequencing depth (coverage) and the length of contig *i*, respectively, and *depth_g_* is the estimated coverage of the whole genome. The multiplicity of contig *i (mul_i_)* is determined by

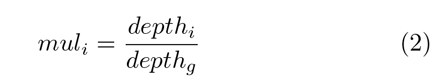

npScarf considers a contig unique if its multiplicity is less than 1.5.

### Bridging unique contigs and filling gaps with repetitive contigs

npScarf next builds the backbone of the genome from the unique contigs. It identifies the long reads that align to two unique contigs, thereby establishing the relative position (*i.e.,* distance and orientation) of these contigs. In order to minimise the effect of false positives that can arise from aligning noisy long reads, npScarf groups reads that consistently support a particular relative position into a bridge and assigns the bridge a score based on the number of supporting reads and the alignment quality of these reads. When two unique contigs are connected by a bridge, they are merged into one larger unique contig. npScarf uses a greedy strategy based on the Kruskal’s algorithm (Kruskal, 1956) that merges contigs from the highest scoring bridges. In the newly created contig, the gap is temporarily filled with the consensus sequence of the reads forming the bridge. npScarf then identifies repetitive contigs that are aligned to this consensus sequence, and use these contigs to fill in the gap.

### Real-time processing

To support real-time analysis of nanopore sequencing, the previously described algorithm can be augmented to process long read data directly from a stream (See Figure 1). In this mode, npScarf employs a mapping method that supports streaming processing such as BWA-MEM (Li, 2013) to aligns each long read to the existing assembly as the read arrives. If the read is aligned to two unique contigs, it is added to the bridge connecting the two contigs. Once the bridge reaches a pre-defined scoring threshold, the two contigs are merged and the gap is filled as above. In case this merging contradicts with the existing assembly, such as the relative distance and/or orientation implied by the bridge are inconsistent with that of the previous used bridges, npScarf revisits the previous bridges to break the smallest scoring contradicting bridge and uses the current bridging instead. The algorithm hence gradually improves the completeness and the quality of the assembly as more data are received.

### Completing bacterial assemblies

We assessed the performance of our algorithm on scaffolding and completing the Illumina assemblies of two bacterial *Klebsiella pneumoniae* strains, ATCC BAA-2146 (NDM-1 positive resistant strain) and ATCC 13883 (type strain). We first sequenced the genomes of these strains with the Illumina MiSeq platform to a coverage of 250-fold, and assembled them with SPAdes (Bankevich et al., 2012) (See Methods). This resulted in assemblies of 90 and 69 contigs that are 500bps or longer, respectively. The N50 statistics of the two assemblies were 288Kb and 302Kb, respectively. We then sequenced the two strains with Oxford Nanopore MinlON using chemistry R7. For ATCC BAA-2146 strain, we obtained 185Mb of sequencing data (~33-fold coverage of the genome), in which 27Mb were 2D (two-directional) reads. The run for strain ATCC 13883 yielded only 13.5Mb of sequencing data (~2.4-fold coverage). We re-sequenced this strain with the improved chemistry R7.3. By combining sequencing data from both experiments for this strain, we obtained a total of 100Mb (~18-fold coverage) data, including 22.5Mb of 2D reads. The quality of the data, described in (Cao et al., 2015), was broadly similar to that reported by other MinION users (Ashton et al., 2015;Jain et al., 2015;Loman and Quinlan, 2014).

As the pipeline was developed after we performed the MinION sequencing runs, we tested our streaming analysis by rerunning the base-calling using Metrichor service. Sequence reads in fast5 format were written to disk, and were instantaneously picked up and streamed to the pipeline by npReader (Cao et al., 2016). In essence, the scaffolding pipeline received sequence data in fastq format in a streaming fashion as if a MinION run was in progress.

During the analysis, the pipeline continuously reported the assemblies’ statistics (the numbers of contigs and the N50 statistic), allowing us to track the completeness of the assembly, as well as the number of circular sequences in the genome. This is especially important for analysis of bacterial genomes where chromosomes and plasmids are usually circular. To validate the resulting assemblies, we compared them with the reference genomes of these strains obtained from NCBI (GenBank Accessions GCA_000364385.2 and GCA_000742135.1). We also ascertained the predicted plasmids in these assemblies by looking for the existence of plasmid origins of replication sequences from PlasmidFinder database (Carattoli et al., 2014).

Figure 2a) and 2b) and present the progress of assembly completion against the coverage of MinION data during scaffolding. As expected, the N50 statistics increased and the number of contigs decreased with more MinION data. We found that for *K. pneumoniae* ATCC BAA-2146 strain, our algorithm required only 20-fold coverage of sequence data (<120Mb) to complete the genome, reducing the assembly to the limit of 5 contigs (one chromosome and four plasmids). Those five contigs were circularised, indicating they were completed. We found these five contigs were in total agreement with the complete genome assembly of the strain, previously sequenced with PacBio and Illumina (Hudson et al., 2014) (See Table 1 and Supplementary Figure 1).

With 18-fold coverage of the MinION data for the *K. pneumoniae* strain ATCC 13883, the assembly was improved to four contigs, in which one was reported to be circular (Contig 4). These contigs were aligned to the reference genome for this strain, which contained 16 contigs in five scaffolds. We found Contig 1 and Contig 2 from the npScarf’s assembly were aligned to the reference scaffold KN046818.1, while Contig 3 and Contig 4 were aligned to two reference scaffolds (See (Table 1 and Supplementary Figure 2). The alignments contained forward and reverse matches. We found the breakpoints of these matches corresponded to the contig joints in the reference scaffolds, indicating the incorrect orientation of contigs in the reference scaffolds. The reference scaffold KN046818.1 size was 5.2Mb suggesting this scaffold was the chromosome and was fragmented into two contigs in the npScarf’s assembly. In examining this chromosomal sequence, we found the two contigs were separated by an rRNA operon of length 7kb. BLAST search revealed the structure of this operon with rRNA 5S, 23S and 16S as the main components. This rRNA operon sequence was also found to be present at five other loci in the genome, which were all resolved. However, there was not any long MinION read aligning to this particular position possibly because of the low yield of this dataset, causing the chromosome sequence to be fragmented. We anticipate this could be resolved with more nanopore sequencing data. Contig 3 (139kb) and Contig 4 (119kb) contained several origin of replication sequences (See (Table 1), suggesting they were plasmid sequences and also Contig 4 was reported to be a circular sequence. In Contig 4, we noticed an extra plasmid origin of replication sequence (ColRNAI) that was not found in the reference genomes (see Table 1). In examining the position of ColRNAI, we found it was in one of the gaps in the reference scaffold, hence not reported in reference assembly.

**Figure. 2:**
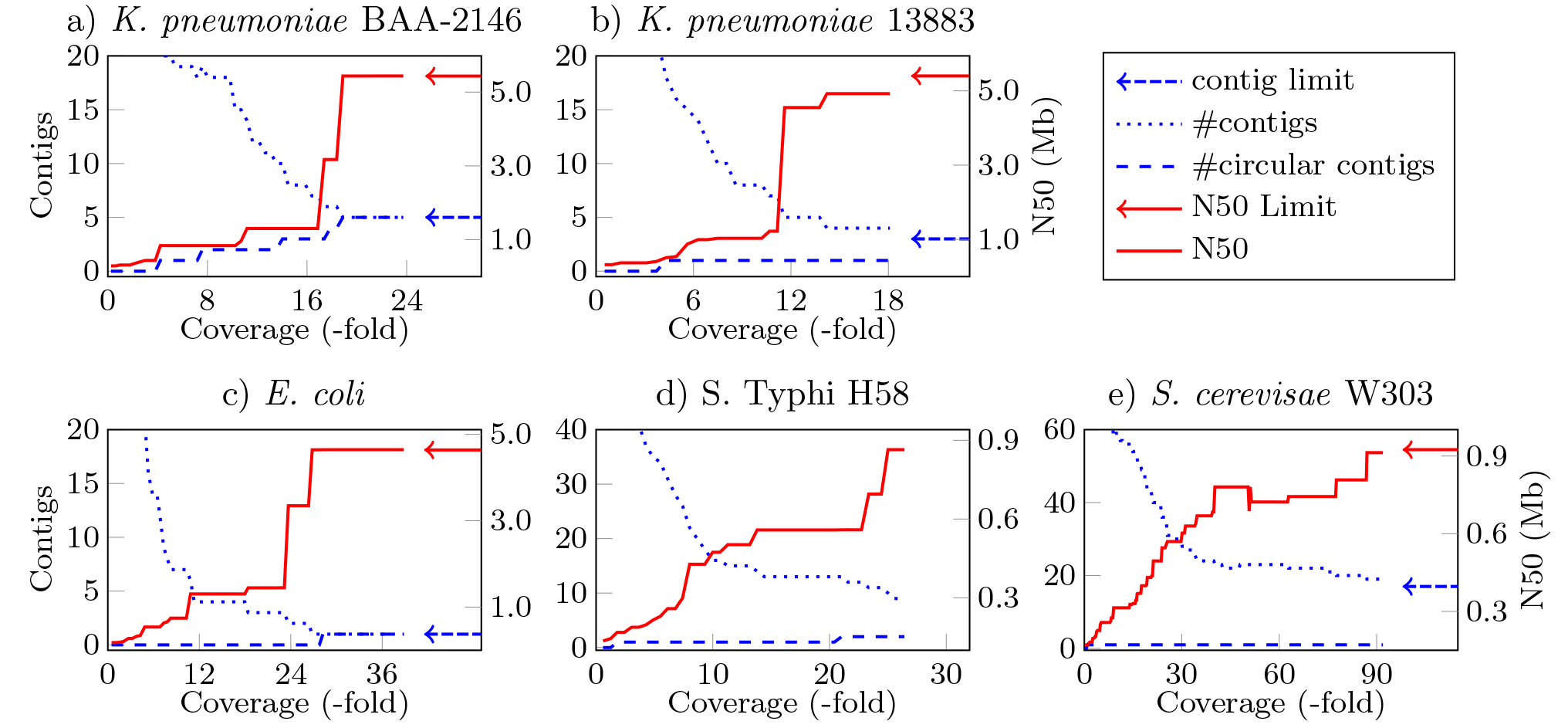
Assembly statistics during real-time scaffolding

**Table 1.**
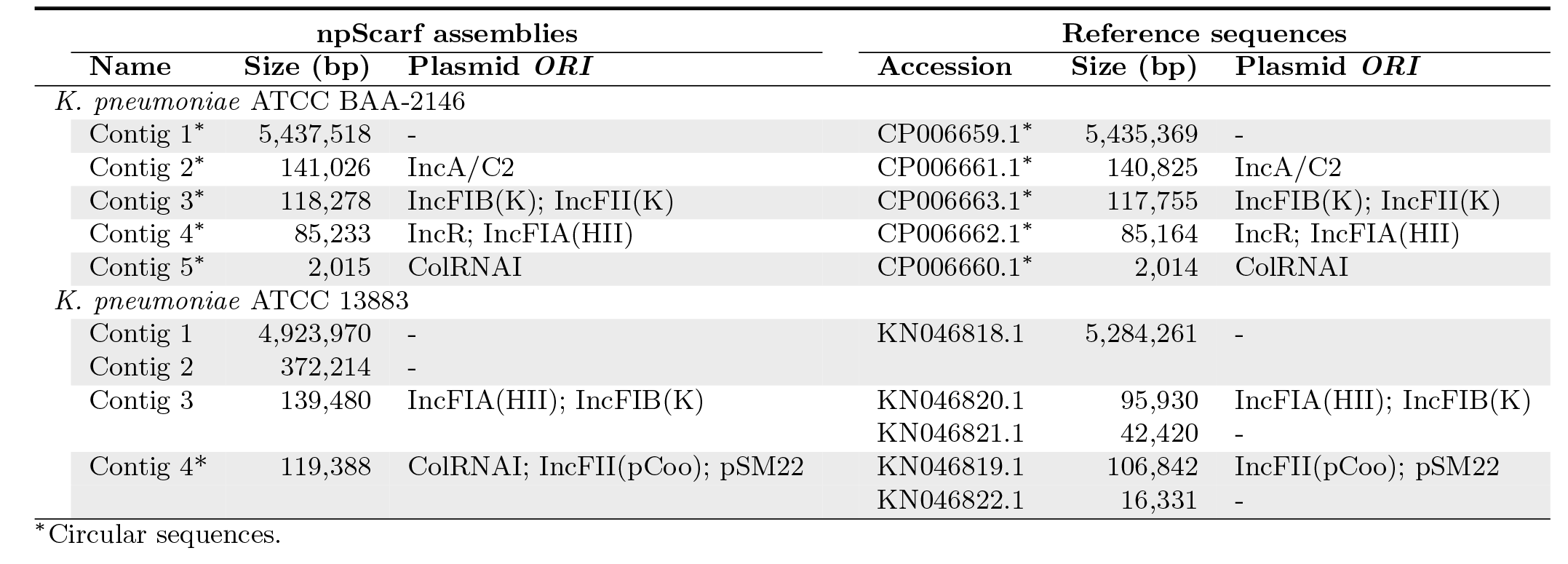
Comparison between the npScarf’s assemblies to the reference genomes of the two*K. pneumoniae*samples

## Real-time identification and plasmids and genomic islands

The ability to complete genome assemblies in streaming fashion also enables real-time analyses that rely on positional information. Such analyses include identifying genes encoded in bacterial genomic islands and plasmids. These functional regions in the bacterial genomes can be horizontally transfered between organisms which is one of the main mechanisms for acquiring antibiotic resistance in pathogenic bacteria. Here we demonstrate these analyses on the multi-drug resistance *K. pneumoniae* ATCC BAA-2146 sample.

Prior to scaffolding the Illumina assembly of the sample, we annotated the assembly using Prokka (Seemann, 2014) to identify the positions of genes and insertion sequences in the assembly. Bacterial genomic islands are genomic regions longer than 8Kb, containing certain classes of genes such as antibiotic resistance genes. In addition, they often carry mobility genes such as transposase, integrase and insertion sequences (IS) (Langille et al., 2010). These sequences generally appear multiple times in the genomes (repetitive sequences), causing genomic islands fragmented in the short read assembly. We ran Island (Mantri and Williams, 2004) and PHAST (Zhou et al., 2011) on the Illumina assembly which together detected six genomic islands. In the annotation, we also found 28 insertion sequences, 14 of them were within 3Kb of the contig ends, suggesting any genomic islands flanked by these insertion sequences were fragmented. During scaffolding of the assembly with nanopore sequencing data, npScarf constructed further four genomic islands which were not previously reported by Island and PHAST (data not shown). Figure 3 presents the structure of such a genomic island, namely Kpn23SapB, and the timeline of its reconstruction. The genomic island harboured three antibiotic resistance genes, aadA (mediates resistance to streptomycin and spectinomycin), sulI (sulfonamides) and ebr (ethidium bromide and quaternary ammonium). The genomic island also carried two copies of the insertion sequence IS26 that flanked the antibiotic resistance genes, and a copy of the insertion sequence IS6100. The presence of these repetitive sequences caused the island to be fragmented into 10 contigs in the Illumina assembly; the three resistance genes were in two different contigs. npScarf required 64.59Mb of data (14-fold coverage of the genome) to report the full structure of the island.

**Figure. 3:**
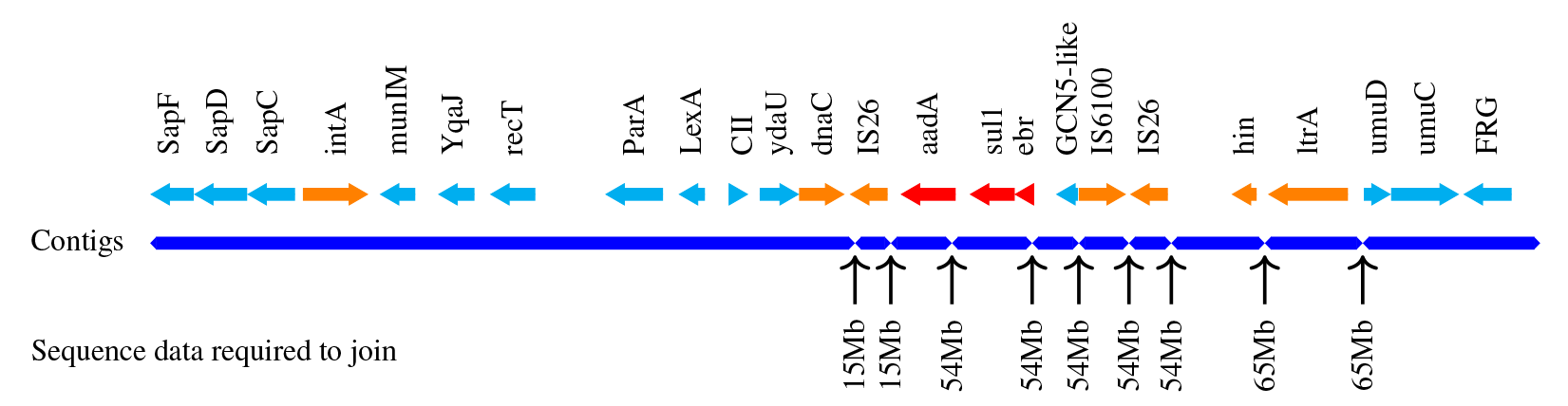
Structure of a genomic island harbouring three antibiotic resistance genes strep, sulI and ebr, anked by mobility genes integrase (int), inverstase (hin), DNA replication (dnaC), and insertion sequences (IS26 and IS6100). The genomic island was fragmentedinto 10 contigs in the Illumina assembly, and was completely resolved with 65Mb out ofthe total of 185Mb of nanopore sequence data.

For real-time detection of plasmid-encoded genes, we identified plasmid origin of replication sequences from the Illumina assembly using the PlasmidFinder database (Carattoli et al., 2014). Contigs that contained a plasmid origin of replication sequence were considered part of a plasmid. Essentially, only 166 genes contained within these contigs couTld be ascertained as plasmid-encoded genes from the Illumina sequencing of the *K. pneumoniae* ATCC BAA-2146 strain. During scaffolding the Illumina assembly, once a contig was added to a plasmid, npScarf reported genes in the contig as plasmid-encode genes. The timelines of detection are presented in the Supplementary Spreadsheet. In particular, we was able to confirm the NDM gene was plasmid-encoded after 46Mb of sequencing.

### Comparison with other methods

We compared the performance of our algorithm against existing methods that were reported to build assemblies with nanopore sequencing. In addition to the two samples presented above, we sourced three other samples reported in the literature including *i.*) an *Escherichia coli* K12 MG1655 strain sequenced to 67-fold coverage with two nanopore R7 flowcells (Quick et al., 2014); ii.) a *Salmonella enterica* serovar Typhi (S. Typhi) haplotype, H58 (Ashton et al., 2015) sequenced to 27fold and iii.) a *Saccharomyces cerevisiae* W303 genome (196-fold) (Goodwin et al., 2015). Of the methods selected for comparison, SPAdes-hybrid (Bankevich et al., 2012), SSPACE-LongRead (Boetzer and Pirovano, 2014), LINKS (Warren et al., 2015) and npScarf were scaffolders whereas Nanocorr (Goodwin et al., 2015) and NaS (Madoui et al., 2015) belonged to the error-correction category. We assembled the Illumina data of these samples using SPAdes (Bankevich et al., 2012) before running the scaffolding methods with nanopore data. SPAdes-hybrid was run by incorporating nanopore data into the assembly (with-nanopore option). The two error-correction tools, Nanocorr and NaS were run on the nanopore sequencing data using about 50-fold coverage of Illumina data, as per suggestion of the respective publications. The corrected reads were then assembled using Celera Assembler (Myers et al., 2000). We observed that the quality of the assemblies produced by Celera Assembler were highly sensitive to the parameters specified in the specification file. We therefore ran Celera Assembler for each data set on three specification files provided by the authors of Nas and Nanocorr, and reported here the most complete assembly obtained.

We evaluated the assemblies in terms of both completeness and accuracy. The completeness of an assembly was assessed by the N50 statistics and the number of contigs that were longer than 500bp. To examine the accuracy of an assembly, we compared that with the closest reference genome of the samples in NCBI (See Methods) to obtain the numberof mis-assemblies and the number of mismatches and short indels. During the test, we recorded the CPU times required by these pipelines to produce the assemblies. The runtimes for the scaffolder methods included times for running SPAdes and for scaffolding, while that for the NaS and Nanocorr included correction time and Celera Assembler time. (Table 2 presents the comparison metrics of all assemblies as reported by Quast (Gurevich et al., 2013) as well as their runtimes.

**Table 2.** Comparison of on assemblies by using npScarf and the comparative methods.

| Method | #Contigs<br>(≥ 500bps) | N50<br>(bps) | Mis-assemblies | Error<br>(per 100kb) | Runtimes<br>(CPUhrs) |  |
| --- | --- | --- | --- | --- | --- | --- |
| *K. pneumoniae* ATCC BAA-2146. Nanopore data: 33X coverage |  |  |  |  |  |  |
| SPAdes | 90 | 288,316 | 0 | 4.72 | 15.63 |  |
| SPAdes-Hybrid | 17 | 3,075,643 | 1 | 6.61 | 16.07 |  |
| SPAdes + SSPACE | 53 | 400,264 | 4 | 12.73 | 15.63 + | 2.3 |
| SPAdes + LINK | 31 | 553,853 | 5 | 16.05 | 15.63 + | 4.03 |
| SPAdes + npScarf (real-time) | 5 | 5,438,323 | 0 | 20.00 | 15.63 + | 1.6 |
| SPAdes + npScarf (batch) | 5 | 5,437,526 | 0 | 22.76 | 15.63 + | 0.84 |
| NaS + CA | 29 | 344,904 | 15 | 18.89 | 324.35 + | 3.49 |
| Nanocorr + CA | 68 | 139,201 | 8 | 141.32 | 312.64 + | 1.37 |
| *K. pneumoniae* ATCC 13883. Nanopore data: 18X coverage |  |  |  |  |  |  |
| SPAdes | 69 | 301,775 | 5 | 6.22 | 16.95 |  |
| SPAdes-Hybrid | 15 | 728,705 | 19 | 8.02 | 16.97 |  |
| SPAdes + SSPACE | 36 | 685,344 | 13 | 12.39 | 16.95 + | 1.48 |
| SPAdes + LINK | 17 | 1,527,003 | 18 | 16.12 | 16.95 + | 1.12 |
| SPAdes + npScarf (real-time) | 4 | 4,923,746 | 21 | 10.84 | 16.95 + | 0.52 |
| SPAdes + npScarf (batch) | 4 | 4,923,952 | 21 | 10.26 | 16.95 + | 0.45 |
| NaS + CA | 38 | 393,946 | 36 | 10.24 | 192.78 + | 6.92 |
| Nanocorr + CA | 60 | 147,647 | 16 | 118.34 | 161.33 + | 2.6 |
| *E. coli* K12 MG1665. Nanopore data: 67X coverage |  |  |  |  |  |  |
| SPAdes | 114 | 176,197 | 0 | 3.51 | 4.38 |  |
| SPAdes-Hybrid | 42 | 4,642,938 | 2 | 1.21 | 4.76 |  |
| SPAdes + SSPACE | 59 | 3,154,619 | 1 | 29.26 | 4.38 + | 3.42 |
| SPAdes + LINK | 50 | 3,317,644 | 2 | 36.19 | 4.38 + | 4.03 |
| SPAdes + npScarf (real-time) | 1 | 4,643,557 | 2 | 13.08 | 4.38 + | 2.43 |
| SPAdes + npScarf (batch) | 1 | 4,645,701 | 2 | 11.72 | 4.38 + | 1.91 |
| NaS + CA | 21 | 873,750 | 19 | 10.60 | 807.19 + | 6.77 |
| Nanocorr + CA | 2 | 4,649,789 | 6 | 10.41 | 213.68 + | 8.49 |
| *S. Typhi* H58. Nanopore data: 26X coverage |  |  |  |  |  |  |
| SPAdes | 89 | 106,832 | 7 | 39.05 | 1.86 |  |
| SPAdes-Hybrid | 27 | 443,374 | 12 | 55.46 | 2.06 |  |
| SPAdes + SSPACE | 34 | 358,489 | 10 | 59.39 | 1.86 + | 1.55 |
| SPAdes + LINK | 20 | 473,170 | 13 | 66.65 | 1.86 + | 1.28 |
| SPAdes + npScarf (real-time) | 9 | 864,338 | 18 | 53.86 | 1.86 + | 0.93 |
| SPAdes + npScarf (batch) | 8 | 864,241 | 16 | 52.01 | 1.86 + | 0.47 |
| NaS + CA | 54 | 211,555 | 17 | 58.87 | 248.32 + | 7.21 |
| Nanocorr + CA | 95 | 36,608 | 9 | 973.63 | 199.85 + | 0 |
| *S. cerevisiae* W303. Nanopore data: 196X coverage |  |  |  |  |  |  |
| SPAdes | 364 | 155,423 | 29 | 124.10 | 20.54 |  |
| SPAdes-Hybrid | 240 | 346,297 | 68 | 158.13 | 67.81 |  |
| SPAdes + SSPACE | 263 | 392,096 | 89 | 136.66 | 20.54 + | 31.54 |
| SPAdes + LINK | 161 | 579,611 | 83 | 143.04 | 20.54 + | 26.97 |
| SPAdes + npScarf (real-time) | 19 | 912,664 | 82 | 141.93 | 20.54 + | 21.28 |
| SPAdes + npScarf (batch) | 17 | 924,022 | 79 | 141.01 | 20.54 + | 18.84 |
| NaS + CA | 121 | 154,851 | 123 | 140.08 | 9811.88 + | 140.69 |
| Nanocorr + CA | 108 | 599,597 | 133 | 197.00 | 7208.08 + | 272.86 |

We ran npScarf in real-time mode, in which nanopore sequencing data are streamed to the pipeline in the exact order they were generated. This allowed us to assess the completeness of the assemblies against the amount of data generated. Figure 2 shows the progress of completing the assemblies for all five samples. As mentioned previously, npScarf produced a complete and a near-complete assemblies for the two *K. pneumoniae* samples (Figures 2a and 2b) with only under 20-fold coverage of nanopore data. For the *E.coli* sample, npScarf required less than 30-fold coverage nanopore data to complete the genome assembly with one circular contig. npScarf also reduced the S. Typhi assembly to only nine contigs (N50=864kb), which was significantly better than the assembly reported by Ashton et al. (2015) from the same data (34 contigs, N50=319kbs).

As for the *S. cerevisae* W303 genome which contains 16 nuclear chromosomes and one mitochondrial chromosome, npScarf generated an assembly of 19 con-tigs(N50=913Kb), substantially fewer than 108 contigs (N50=600Kb) genereated by the next best method (Nanocorr, see (Table 2). We noticed a drop in N50 statistics at the point where about 50-fold coverage of nanopore data were received (Figure 2e). This was because npScarf encountered contradicting bridges and hence broke the assembly at the lowest scoring bridge in lieu of a higher score one. The N50 was then improved to reach the N50 of 913Kb with 90-fold coverage of nanopore sequencing; the assembly did not change with more data (90-to 196-fold). We examined the assembly by comparing that with the *S. cerevisae* strain S288C reference genome. One of the contigs (Contig 17, length=81kb) was reported to be circular which was completely aligned to the mitochondrial chromosome of the reference genome. Ten chromosomes (II, IV, V, VII, IX, X, XI, XIII, XV and XVI) were completely assembled into individual contigs, and three chromosomes (I, III and VIII) were assembled into two contigs per chromosome (See Supplementary Figure 3). We found a mis-assembly 20 that joined chromosome IV and the start of chromosome XIV into Contig 10. The end of chromosome XIV was also joined with chromosome XII into Contig 2. These mis-assemblies essentially fused these three chromosomes into two contigs.

We reran npScarf on the data sets in batch mode in which the scaffolding was performed with the complete dataset. We found that all five assemblies were slightly more complete than that from the real-time mode. In particular, the *S. cerevisae* W303 assembly was further reduced to 17 contigs as chromosomes I and VIII were resolved into individual contigs (data not shown). In this assembly, 12 out of 17 chromosomes were completely recovered to one contig, one chromosome (XIII) was fragmented into two contigs and three chromosomes were fused into two contigs due to mis-assemblies

In all datasets, npScarf consistently produced the most complete assemblies while its accuracy was among the best. It was the only method that could completely resolve the *K. pneumoniae* ATCC BAA-2146 genome (5 contigs, N50 of 5.4Mb) with no mis-assembly requiring only 20-fold coverage of nanopore data; the second most completed assembly (produced by SPAdes Hybrid) contained 17 contigs and had the N50 of only 3.1Mb despite using 33-fold coverage of nanopore sequence data. On the well studied *E. coli* sample where LINK, Nas and Nanocorr were reported to resolve the whole genome with a larger data set (147-fold coverage) (Warren et al., 2015), none of these methods could produce the same result on the 67-fold coverage data set we tested. npScarf on the other hand, was able to reconstruct the genome into one circular contig with as little as 30-fold coverage of the data. On the S. Typhi data set, npScarf produced assemblies with 9 contigs in real-time mode and with 8 contigs in batch mode (N50=864kb), significantly better than assemblies 55 from other methods (over 20 contigs). Similarly, while the *S. cerevisae* W303 assembly produced by npScarf was near complete and N50 statistics reached the theoretical limit of 924kb, whereas other methods produced over 100 contigs and more mis-assemblies or errors.

In terms of running times, we observed that the scaffolding methods were much faster than the error correction counterparts. Both NaS and Nanocorr required the alignment of the short reads to the long reads which were computationally expensive. On the other hands, the scaffolding pipelines required 20 CPU-hours or less to build an assembly from short reads, and between a few hours to around 30 hours to scaffold the assembly with long reads. Apart from SPAdes-Hybrid which performed scaffolding as part of assembling short reads, npScarf was the fastest among other scaffolders with consistently requiring much less scaffolding times. Note the times reported in Table 2 were for processing the entire nanopore dataset, whereas npScarf could be terminated early once a desirable assembly is obtained.

## Discussion

The development of high-throughput long read sequencing technologies such as PacBio and nanopore has opened up opportunities for resolving repetitive sequences to assemble complete genomes and to improve existing genome assemblies. However, the relatively high error rates of these technologies pose a challenge to the accurate assembly the genome sequences. It is natural to combine these long and erroneous reads with more accurate and cheaper short read data for assembling genomes (Bashir et al., 2012;Koren et al., 2012). One such hybrid-assembly approach is to correct the long reads, which are then used to assemble the genome (Bashir et al., 2012;Goodwin et al., 2015;Koren et al., 2012;Madoui et al., 2015) with classical assemblers designed for long and accurate reads such as Celera Assembler (Myers et al., 2000). The approach usually requires large amounts of long read data and excessively high computational resources. The second class of hybrid assemblers harness the long spanning reads to guide extension of contigs in the draft genome assemblies. For example, SSPACE-LongRead (Boetzer and Pirovano, 2014) and Cerulean (Deshpande et al., 2013) rely on alignment of long reads to the assembly graph determine the adjacent contigs. LINKS (Warren et al., 2015) uses a k-mer approach which further improves the running time with a small sacrifice of accuracy. These hybrid-assembly methods, especially those in the scaffolding category, provide economical genome finishing pipelines that can produce high quality genome assemblies from small amounts of long read data on modest computing equipments.

The npScarf algorithm presented in this article is similar to these mentioned scaffolders in the sense that npScarf aligns the long reads to the contigs to build a scaffold of the genome. However, our method estimates the copy number of each contig in the genome and constructs the scaffold from non-repetitive contigs while the repetitive contigs are used to fill the gaps in the scaffold. Consequently, npScarf was demonstrated to be able to generate more complete and accurate assemblies than the competitors, while requiring much less data.

One of the main contributions of our algorithm is that it can process data directly from Metrichor base-caller and report the current status of the analysis in real-time. The pipeline hence allows answering the biological problems at hand at the earliest time possible while sequencing is still in progress. Investigators can also assess the progress of the analysis, and terminate the sequencing once an assembly of sufficient quality and completeness is obtained. This enables the generation of sufficient data necessary for the analysis to guarantee the experiment outcomes and at the same time, avoid costly over-sequencing of data.

One limitation of the current approach is that it relies on the cloud-based base-caller, Metrichor, which can introduce a time-lag. In order to circumvent such as Nanocall (David et al., 2016) or DeepNano (Bož et al., 2016) into our pipeline.

The real-time function to complete genomic sequences open the possibility of *in situ* biological analyses (Cao et al., 2015). Certain biological markers of interests may be identified from short read assembly, but their positions in the genome could only be determined by completing 15 the genome assembly with long reads. We have showed that npScarf can facilitate such analyses in real-time by demonstrating the identification of antibiotic resistance genes encoded in genomic islands and plasmids.

## Methods

### Bacterial cultures and DNA extraction

Bacterial strains *K. pneumoniae* ATCC BAA-2146 and ATCC 13883 were obtained from American Type Culture Collection (ATCC, USA). Bacterial cultures were grown overnight from a single colony at 37° C with shaking (180 rpm). Whole cell DNA was extracted from the cultures using the DNeasy Blood and Tissue Kit (QIAGEN©, Cat #69504) according to the bacterial DNA extraction protocol with modified enzymatic lysis pre-treatment.

### Illumina sequencing and assembly

Library preparation was performed using the NexteraXT DNA Sample preparation kit (Illumina) as recommended by the manufacturer. Libraries were sequenced on the MiSeq instrument (Illumina) with 300bp paired end sequencing, to a coverage of over 250-fold.

### MinION sequencing

Library preparation was performed using the Genomic DNA Sequencing kit (Oxford Nanopore) according to the manufacturer’s instruction. For the R7 MinION Flow Cells SQK-MAP-002 sequencing kit was used and for R7.3 MinION Flow Cells SQK-MAP-003 were used according to the manufacturer’s instruction. For each run a new MinION Flow Cell (R7 or R7.3) was used for sequencing. The library mix was loaded onto the MinION Flow Cell and the Genomic DNA 48 hour sequencing protocol was initiated on the MinKNOW software.

### Data collection

MinION data for the *E. coli* sample Loman and Quinlan (2014) were downloaded from the European Nucleotide Archive (ENA) with accession number ERP007108. We used the data from the chemistry R7.3 run (67-fold coverage of the genome from run accession ERR637419) rather than the chemistry R7 reported in work by Goodwin et al. (2015);Madoui et al. (2015);Warren et al. (2015). Illumina MiSeq sequencing data for the sample were also obtained from ENA (assession number ERR654977). Data from both Illumina and MinION sequencing of the S. Typhi strain (Ashton et al., 2015) were collected from ENA accession number ERP008615. The *S. cerevisae* W303 sequencing data were provided by Goodwin et al. (2015) from the website schatzlab.cshl.edu/data/nanocorr/sudheer.zinovyevcurie.com.

### Data processing

Read data from Illumina sequencing were trimmed with *trimmomatic* V0.32 (Bolger et al., 2014) and subsequently assembled using SPAdes V3.5 (Bankevich et al., 2012). SPAdes was run with the recommended parameters (-k 21,33,55,77,99,127 -careful). SPAdes-Hybrid was run with the inclusion of -nanopore option. SSPACE and LINKS were run on the original SPAdes’ assemblies. For SSPACE, we used the parameters reported to work with MinION reads in Karlsson et al. (2015) (-i 70 -a 1500 -g -5000). In case of LINKS, a script was adapted from the example run for *E. coli* to allow 30 iterations of the algorithms being executed for each data set. NaS and Nanocorr were applied to correct nanopore data from the maximum of 50-fold coverage of Illumina data. The corrected long reads were assembled using Celera Assembler version 8.3 with the configuration files provided by the respective publication.

The Illumina assembly of the *K. pneumoniae* ATCC BAA-2146 sample was annotated using Prokka (version 1.12-beta) with the recommended parameters for a *K. pneumoniae* strain. Plasmid origin of replication sequences in both *K. pneumoniae* assemblies were identified by uploading the assembly to the PlasmidFinder database (Carattoli et al., 2014).

### Real-time analyses

In real-time analysis of the Illumina assembly, npScarf aligned incoming long reads using bwa-mem (Li, 2013) with parameters -k11 -W20 -r10 -A1 -B1 -O1 -E1 -L0 -a -Y -K10000 index -. The -K10000 parameter allowed the alignments to be streamed to the scaffolding algorithm after several reads were aligned.

### Comparative metrics

The assemblies produced by the mentioned methods was evaluated using Quast (V3.2) to compare with the respective reference sequences. The number of contigs, N50 statistic and the number of mis-assemblies were as per Quast reports, while the error rates were computed from sum of number of mismatches and the indel length. The CPU time of each pipeline was measured with the Linux time command (/usr/bin/time -v), the sum of user time 100 and system time was reported. When a pipeline was distributed across a computing cluster, its CPU time was the sum of that from all the jobs.

## Data access

The MinION sequencing data for the two *K. pneumoniae* samples were deposited to ENA under accessions ERR868296 and ERR868298. The MiSeq sequencing data are in the process of depositing to ENA.

## Software Availability

The software presented in this article and its documentation is publicly available at github.com/mdcao/npScarf.

## Author’s contributions

MDC and LC conceived the study. SN, MDC and LC designed and implemented the algorithm. AE performed the bacterial cultures and DNA extractions. DG performed the MinION sequencing and Illumina sequencing. SN and MDC performed the analysis and wrote the first draft of the manuscript. All authors contributed to editing the 20 final manuscript.

## Acknowledgements

MAC is an NHMRC Principal Research Fellow (APP1059354). LC is an ARC Future Fellow (FT110100972). The research is supported by funding from the Institute for Molecular Bioscience Centre for Superbugs Solutions (610246).

